# Noradrenergic input from nucleus of the solitary tract regulates parabrachial activity in mice

**DOI:** 10.1101/2022.09.29.510103

**Authors:** Yadong Ji, Chimdiya Onwukwe, Jesse Smith, Hanna Laub, Luca Posa, Asaf Keller, Radi Masri, Nathan Cramer

**Author notes:** Corresponding authors: Nathan Cramer, Radi Masri. The first two authors contributed equally to this study. The authors declare that they have no competing financial interests.

## Abstract

The parabrachial complex (PB) is critically involved in aversive processes, and chronic pain is associated with amplified activity of PB neurons in rodent models of neuropathic pain. Here we demonstrate that catecholaminergic input from the caudal nucleus of the solitary tract (cNTS_cat_)—a stress responsive region that integrates intero- and exteroceptive signals—causes amplification of PB activity and their sensory afferents. We used a virally mediated expression of a norepinephrine (NE) sensor, NE2h, fiber photometry, and extracellular recordings in anesthetized mice to show that noxious mechanical and thermal stimuli activate cNTS neurons. These stimuli also produce prolonged NE transients in PB that far outlast the noxious stimuli. Similar NE transients can be evoked by focal electrical stimulation of cNTS, a region that contains the noradrenergic A2 cell group that projects densely upon PB. In vitro, optical stimulation of cNTS_cat_ terminals depolarized PB neurons and caused a prolonged increase the frequency of excitatory synaptic activity. A dual opsin approach showed that sensory afferents from the caudal spinal trigeminal nucleus are potentiated by cNTS_cat_ terminal activation. This potentiation was coupled with a decrease in the paired pulse ratio, consistent with an cNTS_cat_ - mediated increase in the probability of release at SpVc synapses. Together, these data suggest that A2 neurons of the cNTS generate long lasting NE transients in PB which increase excitability and potentiate responses of PB neurons to sensory inputs. These reveal a mechanism through which stressors from multiple modalities may potentiate the aversiveness of nociceptive stimuli.

**Significance Statement:** Increased excitability of the parabrachial nucleus (PB), a key integrative hub for aversive stimuli, is linked to amplified pain behaviors. We show that prolonged norepinephrine (NE) transients in PB following noxious stimulation in mice. These NE transients potentiate sensory input to PB and arise, at least in part, from catecholaminergic projections from the caudal nucleus of the solitary tract (cNTS_cat_). We propose that activity this cNTS_cat_ to PB pathway may potentiate the aversiveness of pain.

## Introduction

Chronic pain profoundly affects quality of life (Dworkin et al., 2007; O’Connor and Dworkin, 2009; van Hecke et al., 2014), and afflicts over 100 million people, costing up to $650 billion a year in medical treatment and lost productivity in the United States alone (Institute of Medicine (US) Committee on Advancing Pain Research, 2011). Chronic pain is the most common complaint of patients in outpatient clinics (Upshur et al., 2006) and effective therapies remain limited (Meyer-Rosberg et al., 2001; Dworkin et al., 2013).

Attempts to treat pain have often ignore the role of cognitive, attentional, and emotional aspects of pain perception (Bushnell et al., 2013). While animal models have provided indirect evidence, studies in human participants directly demonstrate the ability of emotional and attentional factors to modify pain perception (deCharms et al., 2005; Loggia et al., 2008; Roy et al., 2008, 2011). There is growing evidence that therapies targeting the motivational-cognitive dimensions of pain may be more promising (Auvray et al., 2010).

A key structure for encoding the affective components of multiple modalities is the parabrachial complex (PB). PB regulates satiety and appetite, sleep and arousal, cardiovascular function, and fluid homeostasis (Hajnal et al., 2009; Martelli et al., 2013; Davern, 2014). PB neurons also encode and coordinate behavioral responses to a range of aversive signals, including itch, cachexia, hypercapnia, and visceral malaise, (Campos et al., 2018; Palmiter, 2018; Chiang et al., 2019) and plays a central role in processing acute and chronic pain, (Gauriau and Bernard, 2002; Roeder et al., 2016, 2016; Uddin et al., 2018; Chiang et al., 2019; Raver et al., 2020). Indeed, in multiple models of chronic pain, PB neurons are hyper-responsive and generate amplified neuronal discharges (Matsumoto et al., 1996; Uddin et al., 2018). These amplified discharges are causally related to the expression of pain behaviors (Asada et al., 1996; Okubo et al., 2013; Raver et al., 2020) and suggest that changes in network excitability and synaptic transmission in PB contribute to chronic pain.

We have shown that synaptic transmission and neuronal excitability in PB are regulated by opioid, cannabinoid and GABA_B_ receptors (Cramer et al., 2021). An additional, and underexplored, source of modulation in PB arises from norepinephrine (NE) and its activation of adrenergic receptors. These receptors are expressed in PB (Unnerstall et al., 1984; Herbert and Flügge, 1995), and manipulation of NE signaling in PB impacts several behaviors, including food intake (Roman et al., 2016; Boccia et al., 2020; Yang et al., 2021), sodium balance (Andrade et al., 2004) and nociception (Wei and Pertovaara, 2006). PB receives dense noradrenergic inputs from several cell groups, but the single largest source originates from the A2 group in the caudal nucleus of the solitary tract (cNTS) (Milner et al., 1986). Neurons in the cNTS are responsive to stress (Rinaman, 2011) and receive exteroceptive input from medullary and spinal dorsal horn neurons (Menetrey and Basbaum, 1987), interoceptive inputs from the vagus and glossopharyngeal nerves, and blood borne signaling molecules (Holt, 2022). Thus, PB and cNTS respond to noxious or aversive stimuli. However, whether catecholaminergic projections from cNTS contribute to NE modulation of synaptic transmission and excitability in PB is unknown.

Here, we tested the hypothesis that noxious stimuli drive NE release in PB by activating catecholaminergic neurons in cNTS (cNTS_cat_), and that this NE release excites PB neurons and potentiates sensory input to PB. Using a combination of *in vivo* fiber photometry, extracellular recordings from cNTS, and *in vitro* optogenetics, we find that the A2 to PB circuit facilitates excitability of PB, and potentiates its sensory inputs.

## Materials and Methods

We adhered to accepted standards for rigorous study design and reporting to maximize the reproducibility and translational potential of our findings as described by Landis et al. (2012) and in ARRIVE (Animal Research: Reporting In Vivo Experiments). We performed an *a priori* power analysis to estimate required sample sizes (Landis et al., 2012). All procedures adhered to Animal Welfare Act regulations, Public Health Service guidelines, and approved by the Institutional Animal Care and Use Committee.

### Animals

We use adult male and female transgenic mice in which Cre recombinase expression is controlled by the tyrosine hydroxylase promotor (TH-Cre) promotor (Savitt et al., 2005). Experimental mice were bred in-house from breeding pairs purchased from the Jackson Laboratory (JAX stock #008601: B6.Cg-7630403G23RikTg(Th-cre)1Tmd/J). We used: 5 male and 3 female mice for *in vivo* fiber photometry recordings from the parabrachial nucleus (PB), 1 male and 2 female mice for *in vivo* stimulation of the caudal nucleus of the solitary tract (cNTS), and 1 male and 4 female mice for *in vitro* recordings.

### Viral construct injection

All viral vector injections were performed in a stereotaxic device under isoflurane anesthesia with Rimadyl for postoperative analgesia. We targeted PB via a small (∼1 to 2 mm) craniotomy at 5.2 mm rostral to bregma, +/-1.5 mm lateral from the midline and injected 500 nL of AAV9-hSyn-NE2h (YL003011-AV9, WZ Biosciences) at a depth of 2.9 mm. The viral construct does not contain a fluorescent reporter, rendering it difficult to differentiate sensor fluorescence from tissue auto-fluorescence. It was also not possible to identify probe tracks, since the relatively brief, acute photometry sessions produced no discernible changes in the tissue. Nevertheless, we have confirmed in dozens of surgeries that the coordinates used reliably target the external PB.

We targeted the A2 region of the cNTS by making a small incision in the meninges over the foramen magnum and injected 500 nL of pAAV5-Syn-FLEX-rc[ChrimsonR-tdTomato] (Addgene # 62723) at a depth of 0.5 mm, 0.2 mm lateral to the midline at the caudal aspect of the obex. A similar approach was used to target SpVc where we injected 500 nL of pAAV5-Syn-Chronos-GFP (Addgene # 59170) at a depth of 0.5 mm, 1.8 mm lateral to the midline. Viral constructs were injected at a rate of 50 nL/min. We waited 3 to 6 weeks for expression prior to recordings.

### Fiber Photometry

Mice were anesthetized with an intraperitoneal injection of 2mg/kg urethane, placed on a stereotaxic frame and a fiber optic probe (400 µM diameter, 0.39NA; RWD Life Sciences) was placed in the right PB at the same coordinates used for NE2h injection (−5.2mm AP, +1.5mm ML, -2.2 to -2.5mm DV). The fiber optic probe was connected to a RZX10 LUX fiber photometry processor running Synapse software (Tucker-Davis Technologies) through a Doric mini cube (Doric Lenses). LEDs at 465 nm (30 μW) and 405 nm (10 μW) were used for NE2h excitation and isosbestic control respectively. LED power was verified and calibrated as needed using a digital optical power meter (Thor Labs). The probes caused minimal tissue damage which was undetectable below the superficial inferior colliculus. Where present, we used this superficial damage to verify our recordings were in the correct rotrocaudal and mediolateral planes above PB.

### Mechanical, thermal and electrical stimulation

We used calibrated forceps to apply 10 consecutive noxious pinch stimuli, ∼ 5 to 10 s duration, to the right (ipsilateral), left (contralateral) hindpaw and tail. Resulting changes in NE2h fluorescence were recorded for at least 60 s and consecutive stimuli were spaced a minimum of 2 minutes apart. We turned the LEDs off between each stimulation to minimize photobleaching of the NE2h sensor.

To generate noxious thermal stimuli, we applied heat stimuli to the dorsocaudal edge of the vibrissae pad and hindlimb foot pad using a laser (AMD Picasso) with the probe (400 μm tip diameter) positioned 5 mm from the skin. The laser power was adjusted with a Jenco Electronics (Grad Prairie, TX) microcomputer thermometer to reach 50°C by the end of a 20 s exposure. We waited at least 3 minutes between consecutive stimuli and inspected the skin after each stimulus for tissue damage, which did not occur.

To electrically stimulate cNTS_cat_, we exposed cNTS and inserted a bipolar stimulating electrode (FHC Inc) to a depth of 0.5 mm and applied trains of electrical stimuli at 5, 10, or 20 Hz (0.2 to 2.5s; 80 to 130 μA). We waited a minimum of 2 minutes between trials. Stimuli were applied bilaterally and, since we did not observe a significant difference between stimulation side, the resulting NE2h response metrics were combined for analysis.

### Anesthetized Electrophysiology Recordings

Animals were anesthetized by intraperitoneal injections of urethane (10% w/v), placed in a stereotaxic frame with a heating pad, and a craniotomy was made over the recording site to target A2 (1-2mm lateral to the obex). Once a responsive cell was identified, either noxious pinch or 50ºC hot water was applied to the hind paw of the animal. Noxious pinch lasted for 1 second and the hot water was applied for 5 seconds. We waited at least 2 minutes between consecutive stimuli and inspected the skin after each stimulus for erythema or tissue damage.

### Brain slice preparation

We anesthetized animals with ketamine/xylazine, removed their brains, and prepared horizontal or coronal slices (300-µm-thick) containing PB, following the method described by (Ting et al., 2014). For recordings, we placed slices in a submersion chamber and continually perfused (2 ml/min) with ACSF containing (in mM): 119 NaCl, 2.5 KCl, 1.2 NaH2PO4, 2.4 NaHCO3, 12.5 glucose, 2 MgSO4·7H2O, and 2 CaCl2·2H2O.

The NE sensor used in this study is an updated version of the NE1 sensor described and validated by Feng et al (Feng et al., 2019). Validation of this second generation, AAV9-hSyn-NE2h (YL003011-AV9, WZ Biosciences), was provided by personal communication (Y. Li, December 5, 2020). As further validation, we obtained brain slices from mice injected with the NE2h viral construct in PB. We recorded changes in sensor fluorescence, as described above for *in vivo* recordings, in response to bath application of NE (2 to 10 μM), 5HT (5 to 50 μM), or NE transport inhibitor (1 nM).

We obtained whole-cell patch-clamp recordings, in voltage-clamp mode, through pipettes containing (in mM): 120 potassium gluconate, 10 potassium chloride, 10 HEPES, 1 magnesium chloride, 2.5 ATP-Mg, 0.5 EGTA, and 0.2 GTP-Tris. Impedance of patch electrodes was 4 – 8 MΩ. Series resistance (<40 MΩ) was monitored throughout the recording, and recordings were discarded if series resistance changed by >20%. All recordings were obtained at room temperature.

To optically activate Chronos or ChrimsonR, we collimated light through a band pass filter and water-immersion 40X microscope objective to achieve whole-field illumination. Light source for each opsin was a single wavelength (470 nm or 550 nm) LED system (CoolLED pE-100, Scientifica, Clarksburg, NJ), controlled through a TTL signal.

### Data Analysis

#### In vivo photometry

We used customized Python scripts adapted from Tucker-Davis Technologies templates and calculated relative changes in fluorescence as outlined in the photometry analysis package, pMat, developed by the Barker lab (Bruno et al., 2021). Briefly, the isobestic control fluorescence was scaled and subtracted from the NE2h fluorescence and event related changes in sensor fluorescence were converted ΔF/F using the 5 second window prior to each stimulation (mechanical or thermal) as baseline. The decay in fluorescence from the initial peak had an initial fast component followed by a slower decay back to baseline. The time constants for these decay rates were calculated by fitting single exponential decay functions to each component.

#### In vitro recordings

Changes in spontaneous synaptic activity produced by optical stimulation were calculated by measuring the frequency and amplitude of synaptic events 5 to 10 s prior to stimulation and 15 s immediately following the end of the stimulus train. Changes in optically evoked EPSCs from SpVc afferents were measured by averaging the amplitudes of 10 to 15 responses to single or paired pulse optical stimuli, at 10 to 20 s intervals, before and after cNTS_cat_ stimulation. We used a 250 ms, -5 mV square pulse at the end of each stimulation sweep to monitor access resistance and estimate membrane resistance (Rm). The amplitudes of evoked responses were normalized to changes in Rm by multiplying the average evoked response by the Rm before and after cNTS_cat_ stimulation. Paired pulse ratios (PPR) were calculated by dividing the average amplitude of the EPSC evoked by the second pulse by average of the EPSC amplitude evoked by the first pulse.

#### In vivo recordings

Recordings were sorted using Offline Sorter (Plexon Inc., Dallas, TX) using dual thresholds and principal component analysis. Responses to thermal stimuli were analyzed with custom MATLAB scripts. Significant responses were defined as firing activity exceeding the 95% confidence interval of the pre-stimulus firing rate. Peristimulus time histograms (PSTHs) were generated to analyze responses to repeated stimuli.

#### Statistics

All statistical comparisons were performed using GraphPad Prism software. When data passed normality tests (D’Agostino & Pearson, Anderson-Darling, and Shapiro-Wilk), we used a paired t-test for comparing before and after or within-animal measures. For data that failed the normality test, we used a paired Wilcoxon test. All mice/cells were included in the analysis, except for the paired pulse ratio data. In this comparison, we only included cells in which the amplitude of the optically evoked EPSC was significantly changed by cNTS_cat_ stimulation as determined by a t-test. As described in the Results, this led to the exclusion of 1 of 6 neurons from PPR analysis. In all tests, a p – value less than 0.05 was considered significant. Data in all graphs are shown as mean/median with 95% confidence intervals unless explicitly stated otherwise in the figure legends.

## Results

### Noxious stimuli drive prolonged elevations of norepinephrine in PB

The dense innervation of the parabrachial nucleus (PB) by noradrenergic afferents suggests that norepinephrine (NE) affects signaling in PB neurons. Based on this hypothesis, we tested the prediction that noxious inputs evoke NE release in PB. We used an AAV construct to drive expression of a fluorescent NE sensor (NE2h) in PB and used fiber photometry to record responses to noxious mechanical and thermal stimuli in anesthetized mice (n = 3 female and 5 male). Both stimulus modalities were tested all but one male mouse.

The properties of this class of sensor were previously characterized by the originating laboratory (Feng et al., 2019). We verified the responsiveness of the sensor using fiber photometry in PB in acute brain slices (4 slices from 2 mice). As shown in Figure 1A, increasing concentrations of bath-applied NE caused an increase in NE2h fluorescence which began to reverse upon washout. This reversal of NE2h fluorescence was slowed by application of 1 nM reboxetine, a NE uptake inhibitor. In contrast, the sensor did not respond to serotonin (5 μM, n = 1).

**Figure 1:**
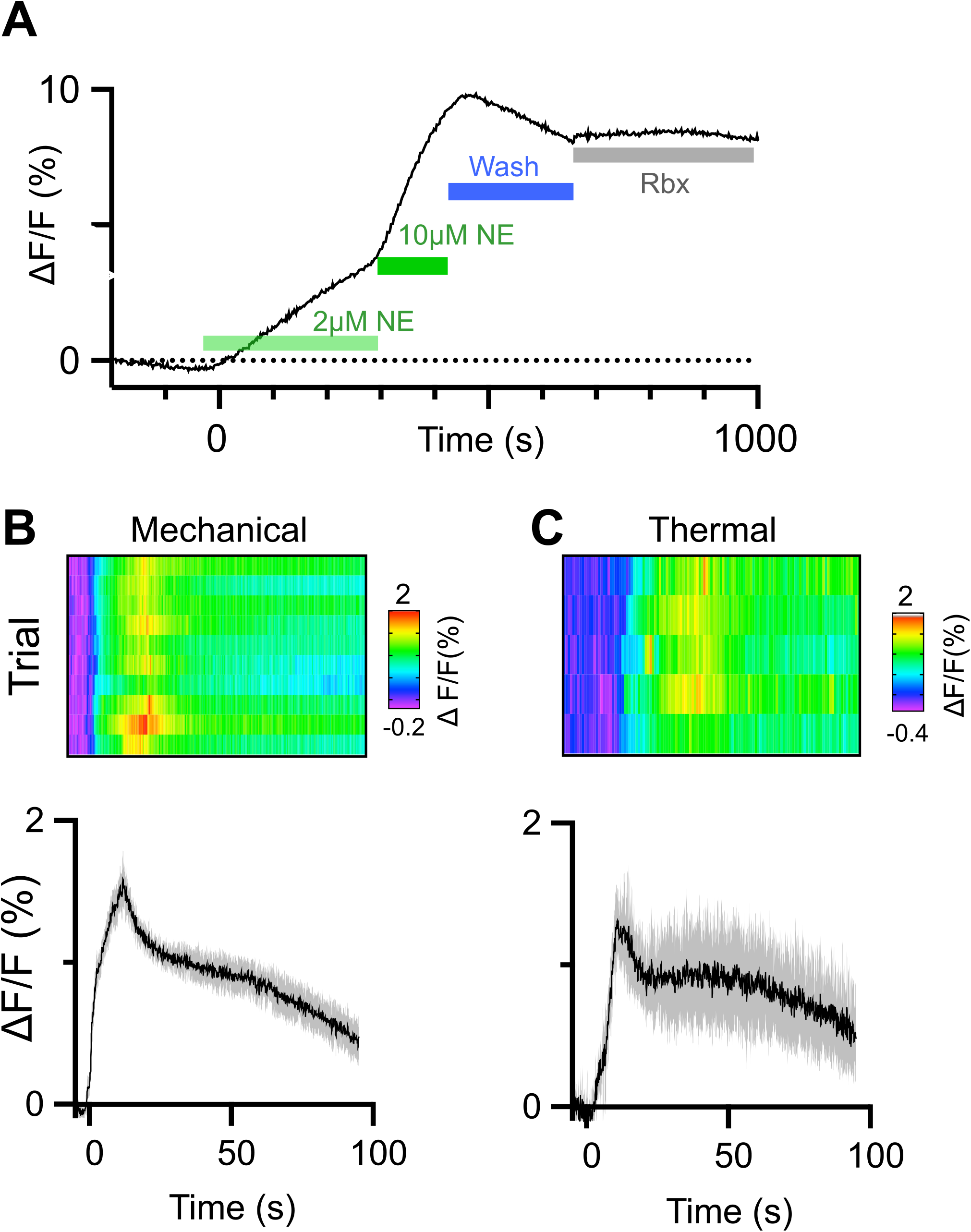
Detecting norepinephrine transients in PB using NE2h sensor. **(A)** Bath application of NE *in vitro* caused a dose dependent increase in NE2h fluorescence in acute PB slices. The decline in fluorescence following washout (blue bar) was partially reversed by the selective NE uptake inhibitor reboxetine (1nM). In anesthetized mice, noxious mechanical **(B)** and thermal **(C)** stimuli reliably produce long lasting NE transients in PB. In each panel, the baseline normalized changes in fluorescence (ΔF/F) for individual trials are shown as heat maps with the mean and 95% confidence intervals of all responses below.

Figures 1 B and C depict examples of averaged NE2h signals and individual trials as heat maps, recorded from PB of the same animal, in response to noxious mechanical stimuli applied to the paw (calibrated pinch; Fig 1B; average of 10 pinches) or thermal stimuli applied to the face (calibrated laser beam; Fig. 1C; average of 5 stimuli). Fluorescence changes were absent in control recordings from 3 mice in which only the reporter (GFP) was expressed. Responses to these noxious stimuli were typified by a rapid increase in fluorescence during the stimulus, followed by a gradual decay in signal intensity towards pre-stimulation values. As analyzed in detail below, the long duration of these responses far outlasted the duration of the noxious stimulus.

#### NE transient magnitude

Fluorescent transients far outlasted the stimulus in all mice tested, despite variation in the maximum amplitude of the response. These features are shown in Figures 2A and B for thermal stimuli applied to the face and hindpaws. The gradient bar in each figure indicates the duration of thermal stimulation. As described in Methods, we used a laser to gradually warm the surface of the skin from ambient temperature to a maximum of 55 °C. Stimuli were applied both contra- and ipsilaterally to the fiber optic probe in PB. Thin lines represent the mean of 5 responses for each mouse while the thick lines with shaded regions indicate the group mean with 95% confidence intervals. To determine if the magnitude of NE2h responses in PB depended on the side of the body stimulated, we compared the area under curve from 0 to 60 s after the stimulus (AUC_0-60s_, Fig. 2C) as well as the peak ΔF/F response (Fig. 2D) obtained from ipsilateral or contralateral stimulation. The mean and 95% CIs and results of paired t-test for each stimulus location are shown in Table 1. Peak ΔF/F values from stimulation of the face was the only metric that showed an effect of laterality, with contralateral stimulation producing a larger response. However, the effect size was small (Cohen’s d = 0.17).

**Figure 2:**
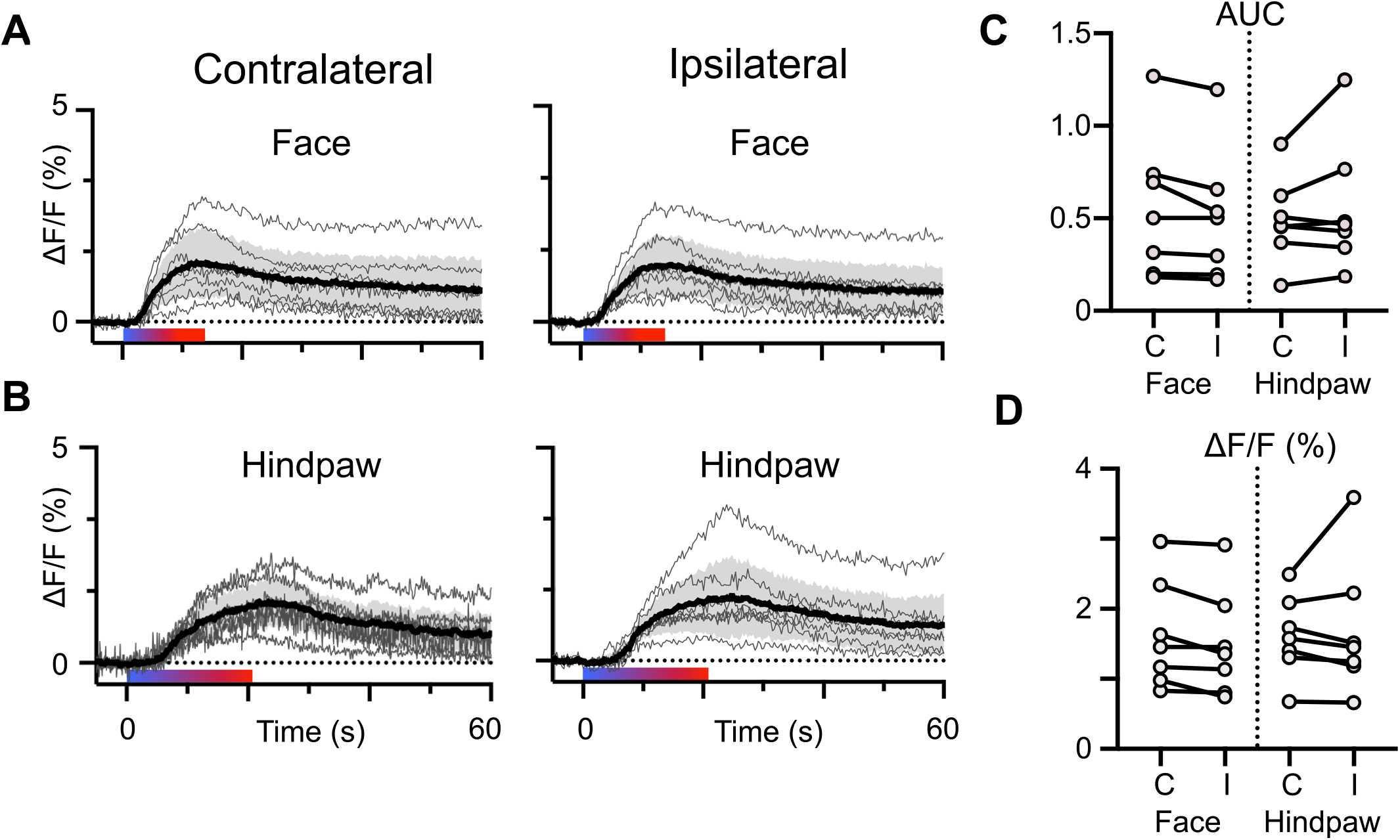
Laterality of noxious heat stimulus has little effect on NE transients. Noxious thermal stimuli applied to the face **(A)** or hindpaw **(B)** produce NE transients that are similar in magnitude and duration whether applied contralaterally or ipsilaterally to the fiber optic probe in PB. In each graph, the light gray traces represent the mean response of an individual mouse to ∼ 5 stimuli. The black trace and shaded region indicate the group mean with 95% confidence intervals. The approximate duration of stimulation across all mice is indicated by the colored bar below each set of traces. The magnitude of transients evoked by contralateral or ipsilateral were not different from each other when measured as the area under the curve **(C)** or peak ΔF/F **(D)**. Paired t-test, p > 0.05.

**Table 1.**

| Table 1: | | AUC | | | $\Delta F/F$ (%) | | |
| --- | --- | --- | --- | --- | --- | --- | --- |
| NE2h response to thermal stimulation |  | Mean | CIs | p-Value | Mean | CIs | p-Value |
| Face | Contra | 0.56 | 0.20 to 0.91 | 0.06 | 1.6 | 0.9 to 2.3 | 0.04 |
|  | Ipsi | 0.51 | 0.18 to 0.84 |  | 1.5 | 0.8 to 2.2 |  |
| Hindpaw | Contra | 0.49 | 0.28 to 0.71 | 0.25 | 1.6 | 1.1 to 2.1 | 0.6 |
|  | Ipsi | 0.56 | 0.24 to 0.88 |  | 1.7 | 0.8 to 2.6 |  |

Mechanical pinch of the left or right hindpaw also reliably produced large and long lasting NE2h transients in PB (Fig. 3A and B). As with thermal responses, the mean responses for each mouse are shown as light grey lines with the group mean and 95% CIs indicated by the bold line and shaded region. Mean AUC_0-60s_ and peak ΔF/F for contra- and ipsilateral stimuli are provided in Table 2. Stimuli applied contralateral to the fiber optic probe generated larger AUC and peak ΔF/F responses than those applied ipsilaterally (Table 2, Cohen’s d = 0.3 and 0.4). These data suggest that, although there is some laterality in the magnitude of NE transients evoked by noxious mechanical stimuli, stimuli of either modality on either side of the body evoke substantial and prolonged NE responses in PB. The similarity in NE2h transients evoked by mechanical pinch of the base of the tail (Figure 4, Table 2), further suggest that the elevations in NE within PB largely reflect the presence of a noxious stimulus rather than its specific location.

**Figure 3:**
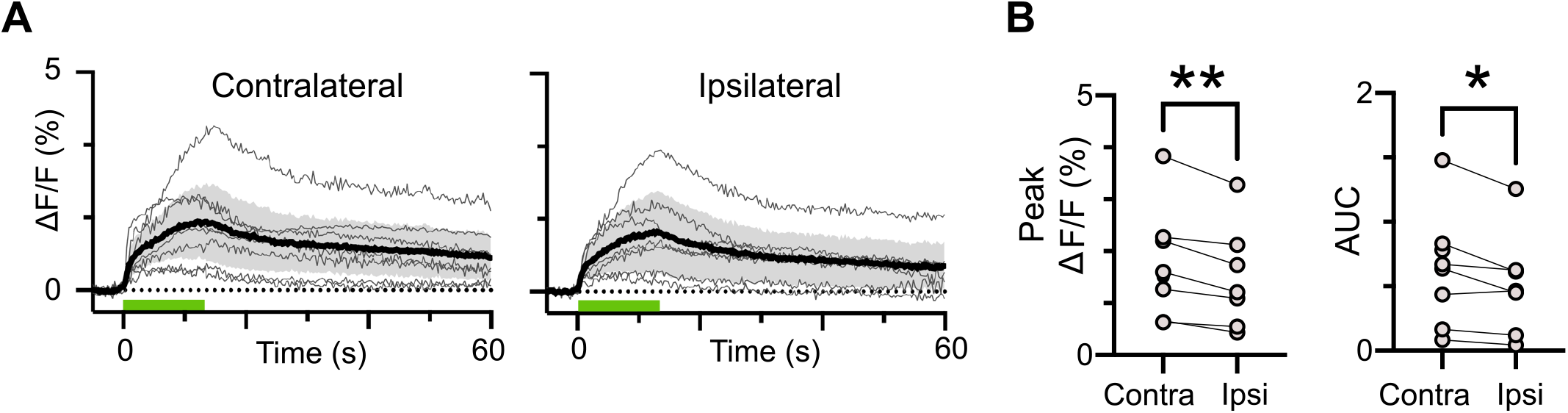
Contralateral mechanical stimuli produce larger magnitude NE transients in PB. **(A)** Responses of individual mice (light traces) and mean with 95% confidence intervals (dark trace with shaded region) to noxious pinch applied to the hindpaws (green bar). Contralateral stimuli produced transients that were greater in magnitude measured as the area under the curve **(B)** and peak ΔF/F. Paired t-test, * = p < 0.05, ** = p < 0.01.

**Figure 4:**
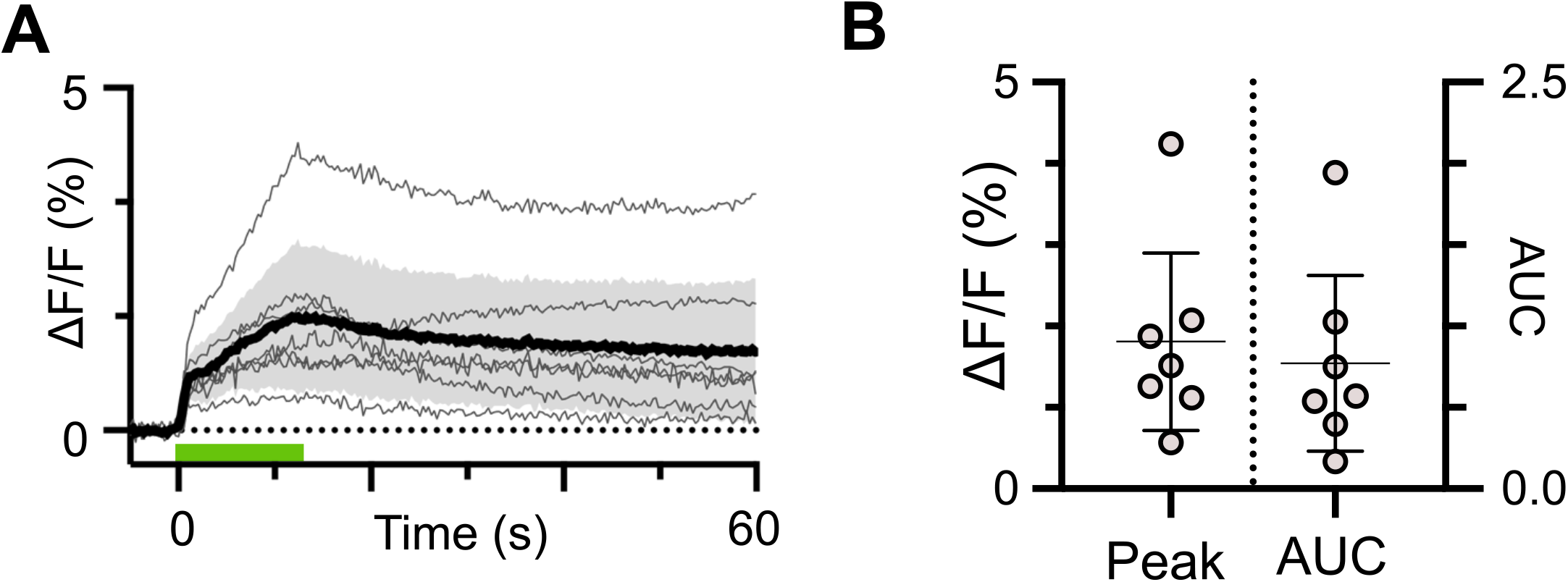
Noxious mechanical stimulation of the tail produces NE transients in PB. **(A)** Responses of individual mice (light traces) and mean with 95% confidence intervals (dark trace with shaded region) to noxious pinch applied to the tail (green bar) and corresponding area under the curve and peak ΔF/F values in **(B)**.

**Table 2.**

| Table 2: | | AUC | | | $\Delta F/F$ (%) | | |
| --- | --- | --- | --- | --- | --- | --- | --- |
| NE2h response to mechanical stimulation |  | Mean | CIs | p-Value | Mean | CIs | p-Value |
| Hindpaw | Contra | 0.61 | 0.18 to 1.0 | 0.06 | 1.8 | 0.7 to 2.8 | 0.06 |
|  | Ipsi | 0.51 | 0.14 to 0.88 |  | 1.5 | 0.5 to 2.4 |  |
| Tail |  | 0.77 | 0.23 to 1.3 | N/A | 1.8 | 0.7 to 2.9 | N/A |

#### NE transient kinetics

To quantify the time course of noxious stimulus-evoked NE transients in PB, we measured onset latencies and time constants of the decay from the peak ΔF/F values towards pre-stimulation levels. Figure 5A and B show the mean responses to all mechanical and thermal stimulation depicted in Figures 2 through 4, but here focus on the time around the stimulus. Latencies for individual mice are shown as group data in Figure 5C and D for mechanical and thermal stimulation, and corresponding means and 95% CIs are shown in Table 3. Although there were differences in onset latencies between stimulus modalities and stimulus locations, the differences likely reflect experimental rather than biological factors. Our mechanical stimulus, noxious pinch, reached the experimental threshold rapidly (< 1s) while the heat stimulus warmed the skin gradually, taking ∼ 20s to reach 50°C. In addition, the thicker skin of the hindpaw is likely less sensitive than the skin on the face near the vibrissae. Thus, the differences in latencies likely reflect the rate at which each modality became noxious. The absence of a NE2h response to low, presumably innocuous temperatures, is consistent with other studies showing NTS neurons preferentially respond to aversive stimuli (Pezzone et al., 1993; Rinaman, 2011).

**Figure 5:**
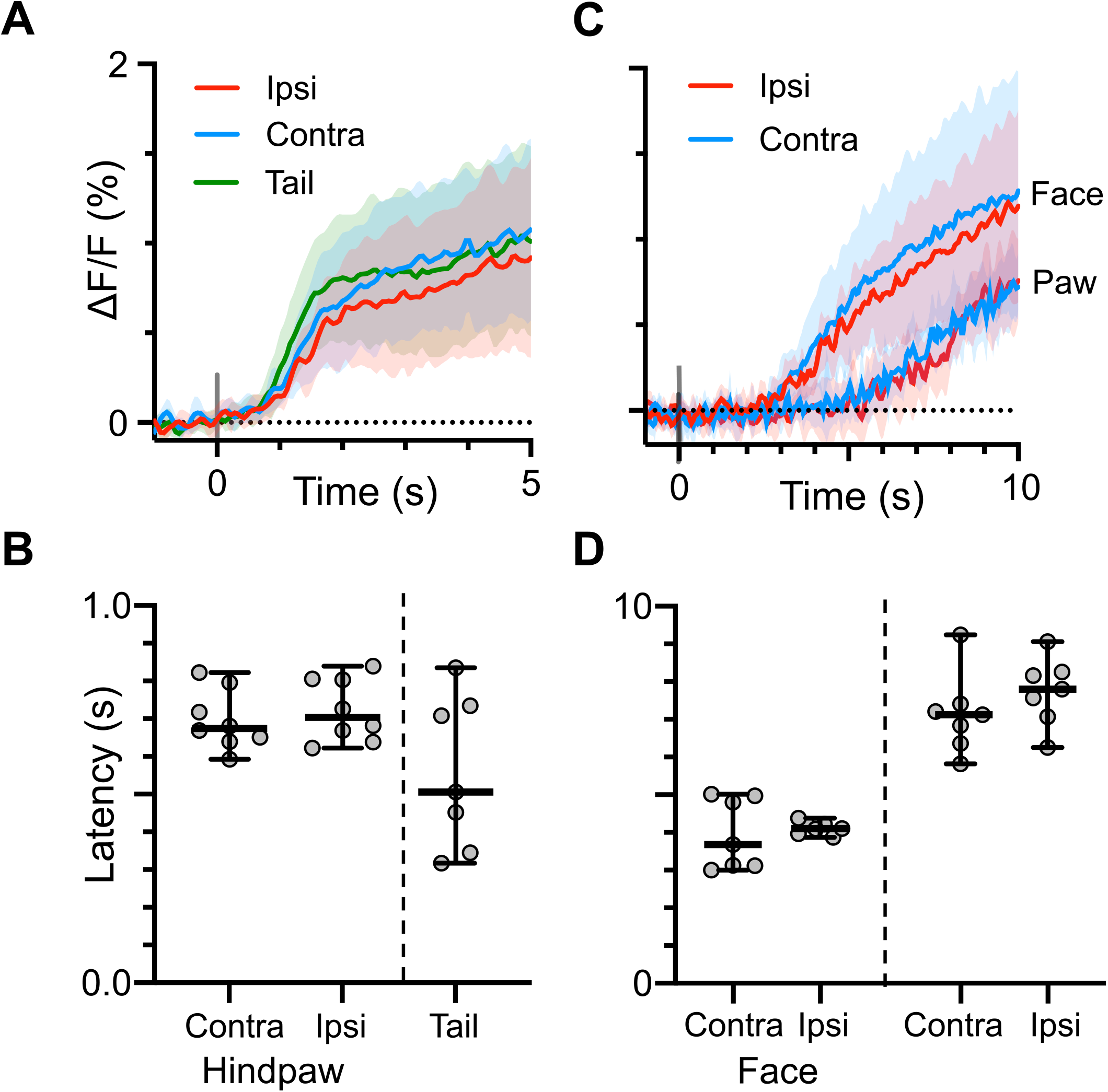
NE transient onset latencies vary by stimulus modality. Expanded views of NE transients evoked by mechanical **(A)** or thermal **(C)** stimulation (group mean and 95% CIs). Latencies from the beginning of the stimulus to onset of the NE2h response were shorter for mechanical stimuli **(B)** and varied by stimulus location for thermal stimuli **(D)**, likely reflecting the rate at which each stimulus became noxious. There were no differences in latencies between contralateral and ipsilateral stimuli of the same modality and location. Paired t-test, p > 0.05.

**Table 3.**

| Table 3:<br>NE2h onset latencies | Mechanical |  |  | Thermal |  |  |  |
| --- | --- | --- | --- | --- | --- | --- | --- |
|  | Hindpaw |  | Tail | Face |  | Hindpaw |  |
|  | Contra | Ipsi |  | Contra | Ipsi | Contra | Ipsi |
| Mean | 0.70 | 0.72 | 0.56 | 4.00 | 4.10 | 7.10 | 7.70 |
| Lower 95% CI | 0.63 | 0.65 | 0.37 | 3.10 | 4.00 | 6.20 | 6.90 |
| Upper 95% CI | 0.76 | 0.79 | 0.74 | 4.80 | 4.30 | 8.10 | 8.60 |

**Table 4.**

| Table 4:<br>NE transient<br>decay rates | Mechanical Decay Rate ($\tau$) | | | Thermal Decay Rate ($\tau$) | | | |
| --- | --- | --- | --- | --- | --- | --- | --- |
|  | Hindpaw |  | Tail | Face |  | Hindpaw |  |
|  | Contra | Ipsi |  | Contra | Ipsi | Contra | Ipsi |
| Mean | 17.15 | 17.43 | 15.58 | 14.85 | 15.62 | 15.42 | 19.86 |
| Lower 95% CI | 12.41 | 13.18 | 8.842 | 10.46 | 11.66 | 11.51 | 13.09 |
| Upper 95% CI | 26.16 | 24.79 | 40 | 23.58 | 22.56 | 22.63 | 38.6 |

Compared to the relatively rapid rise in NE2h signal during the stimulus, the decay in fluorescence occurred more slowly and could remain elevated for over a minute (Figs 2 to 4). We quantified the rate of this decay by fitting a single exponential decay function to the 30 s time window following the peak in the group ΔF/F response. Groups consist of the mean ΔF/F responses from all mice for each stimulus modality and location. Although the stimuli varied in modality, duration and intensity depending on the region of the body being stimulated, the mean rates of decay in peak NE2h fluorescence were similar and had largely overlapping 95% confidence intervals. This suggests that the rate of NE clearance from PB is independent of the type of evoking noxious stimulus used in this study.

### Stimulation of cNTS_cat_ drives prolonged norepinephrine flux in PB

A2 noradrenergic neurons of the caudal nucleus of the solitary tract (cNTS_cat_) are the single largest input of NE to PB. They receive direct excitatory inputs from visceral sensory afferents (Appleyard et al., 2007) and respond to homeostatic challenges, physical and psychological stressors, and peripheral inflammation (Dayas et al., 2001; Hollis et al., 2004). We tested whether these neurons drive NE transients in PB using electrical stimulation of the cNTS in anesthetized mice. We used fiber photometry, as described above, to monitor changes in NE2h fluorescence evoked by 0.2 to 2.5 second stimulus trains applied to cNTS at 5, 10 and 20 Hz (80 to 130 μA). Sample NE2h responses to cNTS stimulation are shown in Figure 6A, and demonstrate that higher stimulation frequencies, at a fixed intensity, produced larger amplitude transients. This result was consistent across 3 mice (Fig. 4B), suggesting that the magnitude of NE transients in PB are proportional to the activity of catecholaminergic neurons of the cNTS. There was no significant difference in the transients evoked by stimulating either the contra– or ipsilateral cNTS (p = 0.7, paired t-test). However, the decay time constant for these transients was similar to that evoked by noxious stimulation (τ = 20, 95% CIs: 13 to 27).

**Figure 6:**
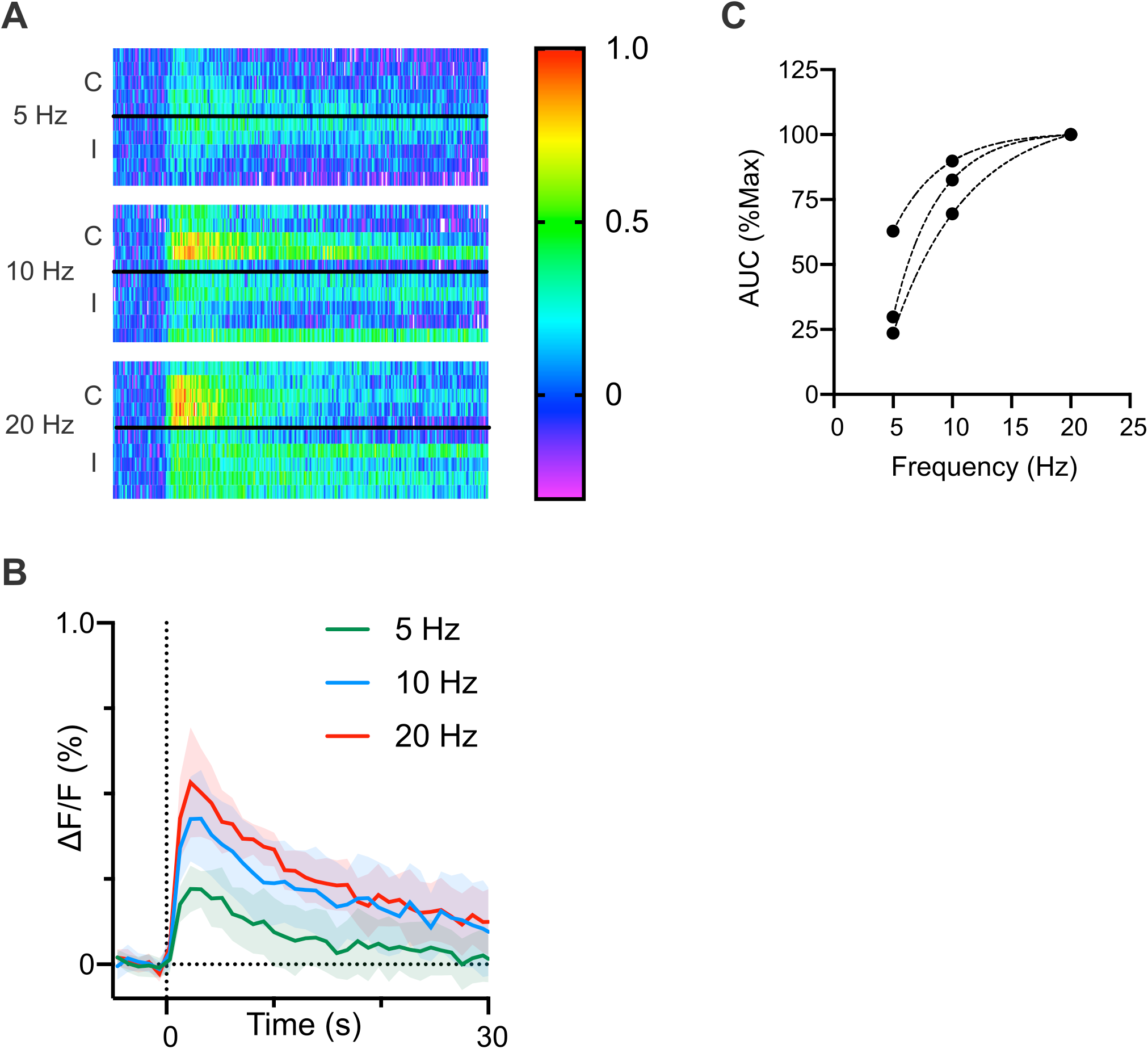
Electrical stimulation of cNTS generates long lasting NE transients in PB. **(A)** Example heat maps from one mouse showing the change in NE2h fluorescence evoked by electrically stimulating cNTS at different frequencies but fixed intensity. The letters “C” and “I” indicate trials where the stimulus was applied to the contralateral and ipsilateral cNTS relative to the fiber photometry probe. **(B)** Mean responses for each stimulus frequency for the data shown in (A). We observed a similar frequency dependent response in 3 mice **(C)**. There was no effect of stimulation side at any frequency, so data from contra- and ipsilateral stimuli were combined for analysis (paired t-test, p > 0.05).

### cNTS neurons respond to noxious stimulation

Our data demonstrate that both noxious stimulation and direct electrical stimulation of cNTS drive prolonged NE transients, supporting the hypothesis that dense NE projections arising from cNTS are a key source of noradrenergic regulation of PB. To investigate this relationship further, we performed single unit recordings from cNTS neurons. Figures 7A and B show representative raster plots and peristimulus time histograms from two cNTS neurons in response to noxious pinch and heat. In both instances, these neurons responded to noxious stimulation with a pronounced increase in firing frequency. We obtained recordings from 9 neurons (4 pinch, 5 thermal) from 9 mice. Based on the baseline firing frequency, one pinch responsive neuron was identified as an outlier (ROUT, Q = 1.0) and removed from further analysis. In the remaining neurons, noxious stimulation increased the median firing frequency from 0.8 Hz (95%CI 0 to 2.5 Hz) to 4.7 Hz (95% CIs: 2.4 to 11 Hz; p = 0.02, Wilcoxon matched pairs test, Figure 7C). As observed with NE transients, the neuronal response was substantially delayed relative to the stimulus onset (1.6 s, 95% CIs: 0.2 to 3.0 s) suggesting that these neurons were not responsive to low intensity mechanical or thermal stimulation. These results demonstrate that at least a subset of neurons in cNTS respond to noxious stimuli.

**Figure 7:**
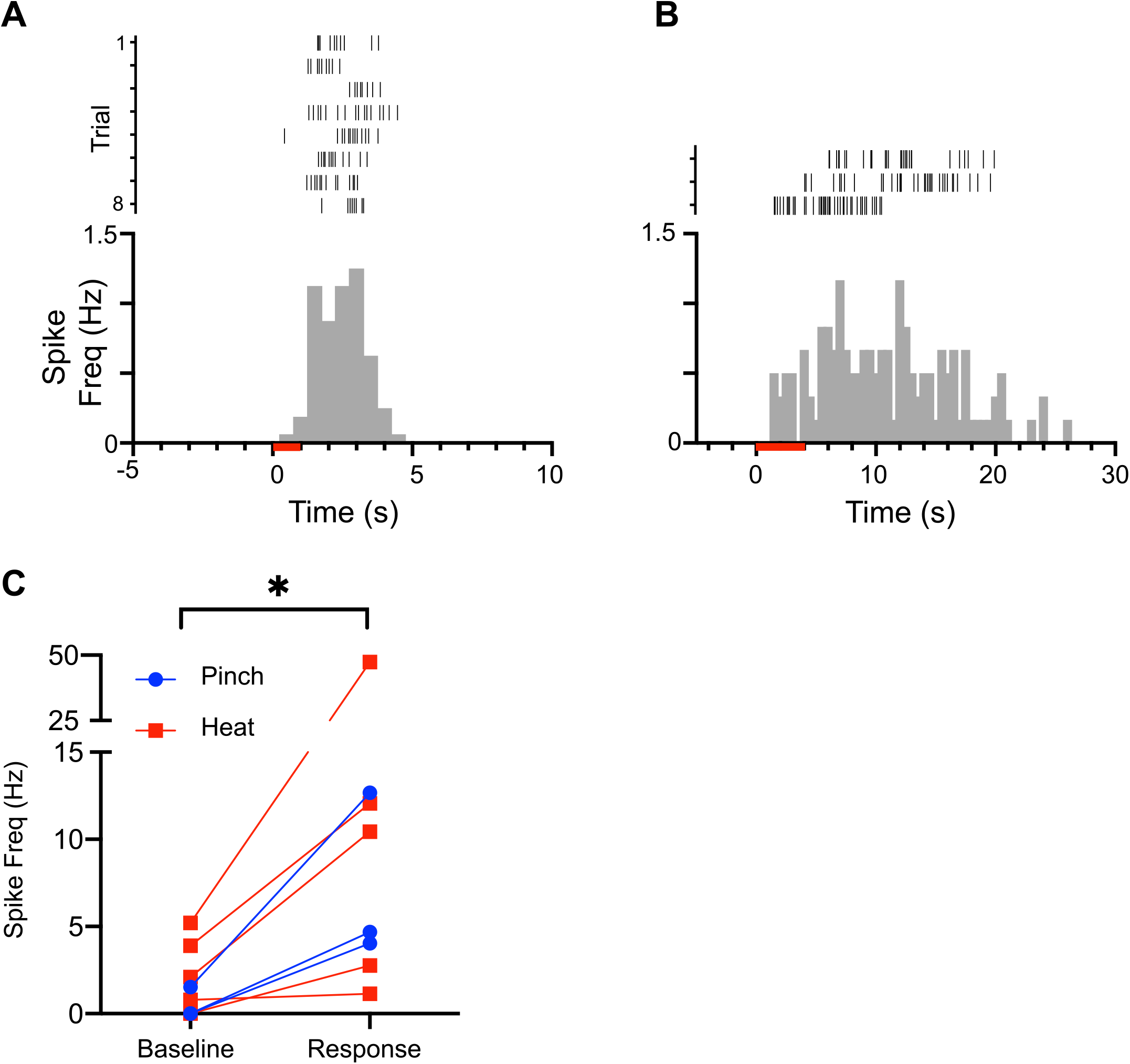
cNTS neurons respond to noxious stimulation. Raster plots and peristimulus time histograms of single unit recordings from unidentified cNTS neurons responding to noxious mechanical **(A)** and thermal **(B)** stimuli. **(C)** Group data from 8 neurons (3 mechanical and 5 thermal) reveal a significant response to noxious stimuli by cNTS neurons (Wilcoxon matched-pairs test, * = p < 0.05).

### cNTS_cat_ Stimulation modulates spontaneous synaptic activity in PB

The observation that NE2h signals remain elevated in PB long after a noxious stimulus suggests that NE afferents may produce long lasting modulation of synaptic transmission in PB. That direct stimulation of the cNTS also evokes robust, prolonged NE transients suggests that A2 neurons may be a prominent source of this modulation. We tested this hypothesis by asking whether activation of cNTS_cat_ inputs has long-term effects on synaptic activity in PB neurons recorded in a slice preparation. To do this, we expressed the red-shifted opsin ChrimsonR (AAV5-syn-Flex-rc(chrimsonR-tdTomato)) in cNTS_cat_ neurons in 5 mice (4 female, 1 male), allowing us to stimulate these afferents in parabrachial slices. We used a train of optical stimuli (10 Hz, 10 s) because sustained depolarization of presynaptic terminals is required for NE release (Schmidt et al., 2019). As shown in the representative voltage clamp recording in Figure 8A, activation of cNTS_cat_ afferents in PB with a train of optical stimuli evoked a short latency, long duration excitatory postsynaptic current in PB neurons. The frequency of synaptic activity also increased during cNTS_cat_ activation and remained elevated after the stimulus ended (Fig. 8B; p = 0.01, paired t-test, Cohen’s d = 1.1, n = 8 neurons from 5 mice) with a mean increase of 370% (95% CIs: 150 to 590 %) relative to the pre-stimulus period (Fig 4B). cNTS_cat_ stimulation did not affect the amplitude of synaptic events (Fig. 8C; mean: 110% of pre-stimulus amplitudes, 95% CI: 90 to 140 %, p = 0.7 paired t-test), despite causing a small reduction in the membrane resistance (Fig. 8D) to an average of 87 % of the starting value (95% CIs: 79 to 96%, p = 0.03, paired t-test, Cohen’s d = -0.2). The decrease in membrane resistance may reflect activation of postsynaptic adrenergic receptors or increase in spontaneous synaptic activity.

**Figure 8:**
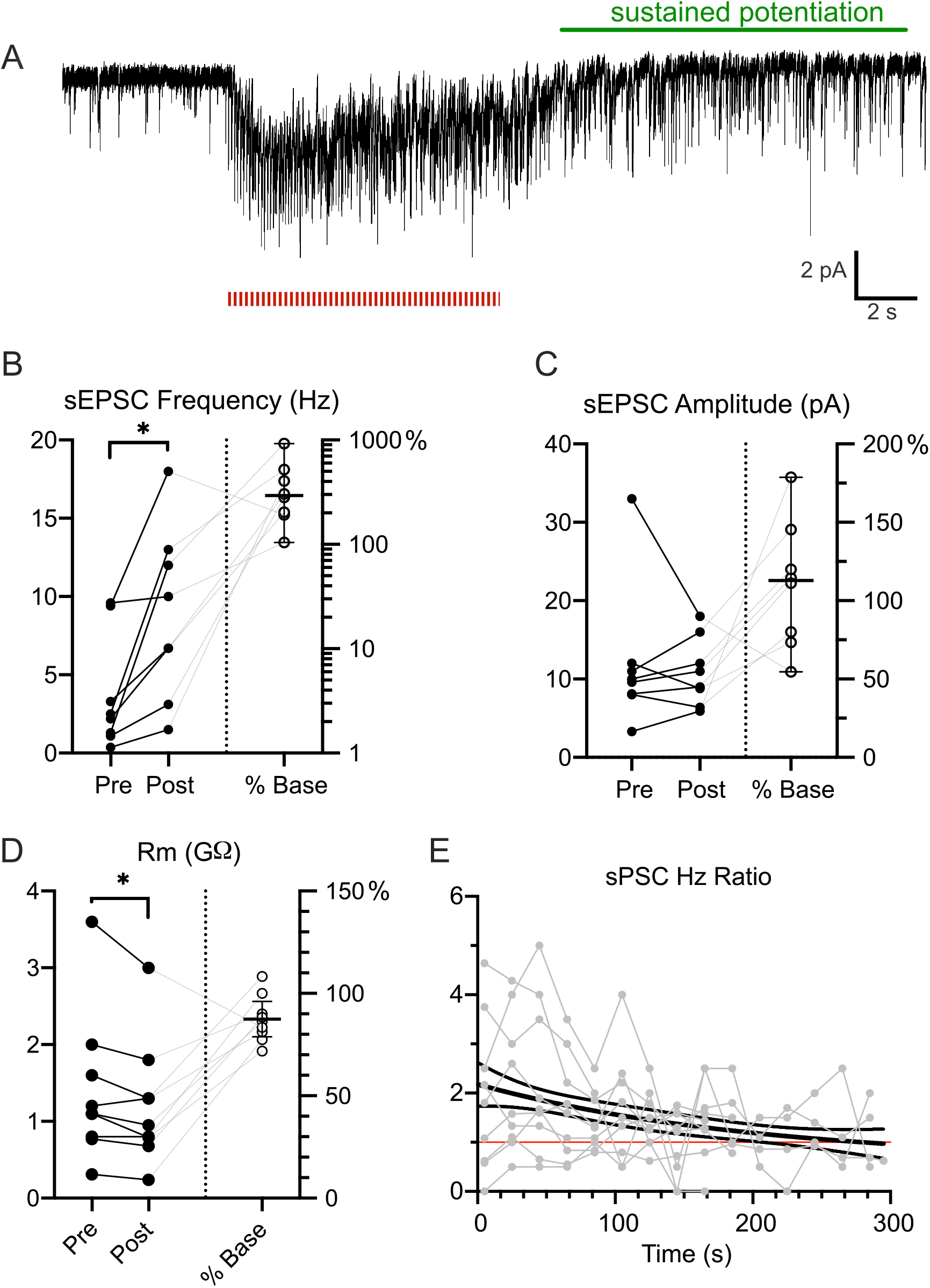
cNTS_cat_ afferent stimulation potentiates excitatory synaptic transmission in PB. **(A)** Representative voltage clamp recording from a PB neuron showing the sustained increase in synaptic activity after A2 afferent stimulation (10 Hz, 10 s, red bars). (**B)** Synaptic activity increased to a mean of 370% of pre-stimulation values in 8 neurons. There was no effect of cNTS_cat_ stimulation on event amplitude **(C)**, despite a small reduction in membrane resistance **(D)**. The cNTS_cat_ -evoked increase in sEPSC frequency was prolonged in 5 neurons and decayed back to pre-stimulus values with a time constant of 100 s.

In a subset of neurons (5 of 8), the increase in synaptic activity remained elevated at least 20 s after the cNTS_cat_ stimulation ended. In these cells, we measured the rate of synaptic events at 5 to 20 second intervals and normalized these values to the median frequency prior to cNTS_cat_ afferent stimulation. Using nonlinear regression, we determined that the decay rate for cNTS_cat_ mediated potentiation to be 100 s (95% CI: 50 to 700 s; Fig. 8E). Further experiments are needed to identify whether this noradrenergic modulation is a general phenomenon within PB or whether specific cell types and/or pathways within PB are targeted.

### cNTS_cat_ stimulation modulates trigeminal afferents in PB

The parabrachial nucleus subserves a range of exteroceptive and interoceptive modalities. We tested whether NE inputs from cNTS_cat_ specifically affect sensory inputs to parabrachial from spinal trigeminal nucleus caudalis (SpVc) using a dual opsin strategy based on (Klapoetke et al., 2014). In TH-Cre mice, we injected pAAV5-Syn-Chronos-GFP into SpVc and pAAV5-Syn-FLEX-rc[ChrimsonR-tdTomato] into cNTS to express Chronos-GFP and CrimsonR-RFP in SpVc and cNTS_cat_ neurons respectively and allowing at least 3 weeks for expression before generating acute brain slices for recording. To ensure we could activate each pathway separately, we used narrow band LEDs as light sources and passed each beam through a band pass filter to minimize the potential for cross-activation of the two opsins. We tested the specificity of this approach explicitly by verifying that each opsin could only be activated when the proper light source was paired with the matching filter. An example of these tests is shown in voltage clamp recordings in Figure 9A. In similar tests across 7 neurons, we did not observe any evidence of cross-activation.

**Figure 9:**
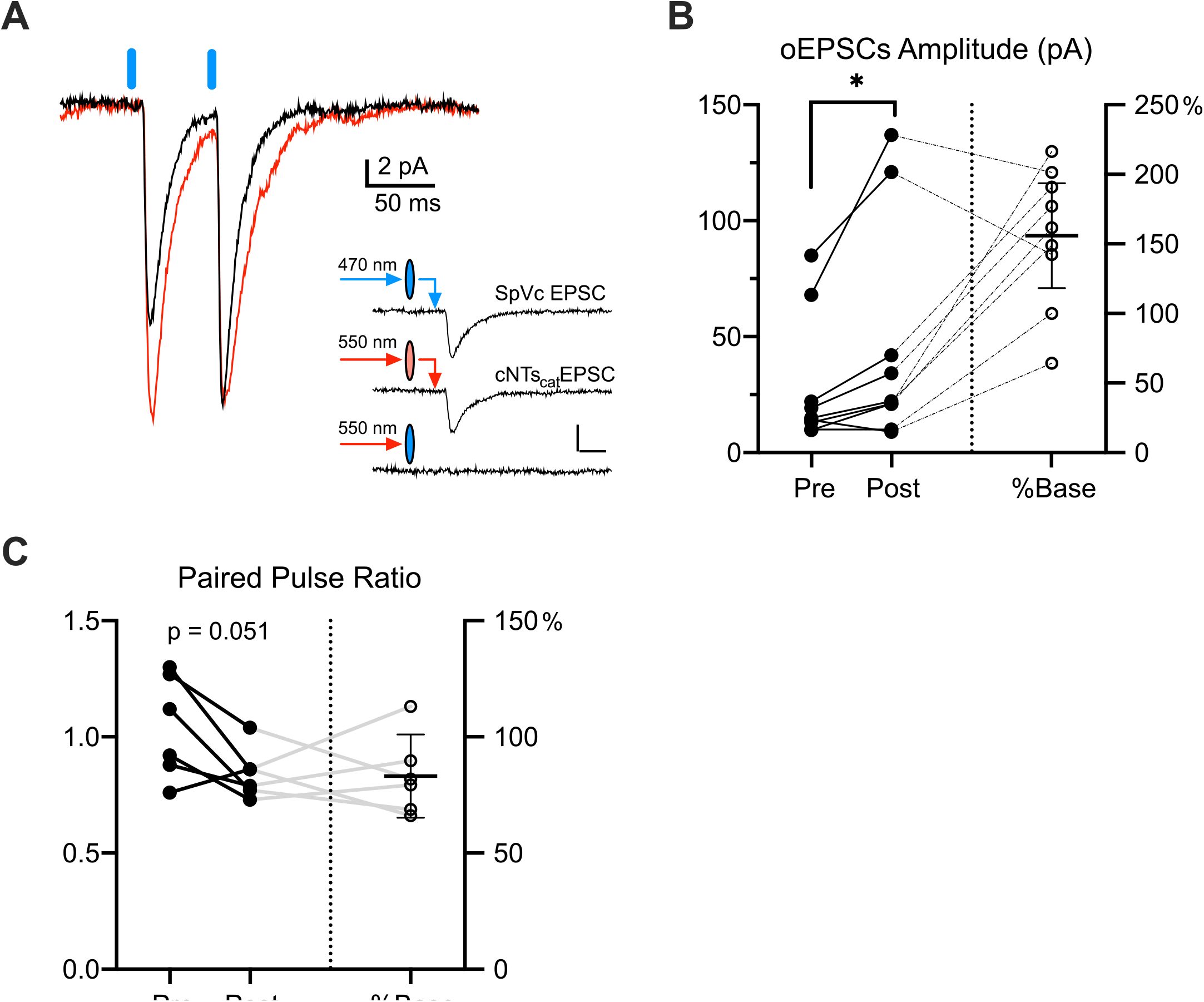
cNTS_cat_ stimulation potentiates sensory inputs to PB neurons. **(A)** Representative EPSCs in PB neurons evoked by paired optical stimulation of SpVc afferent terminals before (black) and after (red) cNTS_cat_ optical afferent stimulation. Inset shows example recording verifying the specificity of optical activation. **(B)** EPSCs evoked were significantly increased by cNTS_cat_ activation by an average of 154 %. **(C)** Paired pulse ratio (PPR) was reduced to 80% of the baseline value (95% CIs: 70 to 100%) at a significance of p = 0.051 (paired t-test, n = 6).

To test whether SpVc inputs to PB could be potentiated by local cNTS_cat_ afferents, we measured the amplitude of SpVc EPSCs (EPSC_SpVc_) evoked by single or paired pulse optical stimuli before and after cNTS_cat_ afferent activation (10 Hz train for 10 s). We obtained recordings from 9 neurons (from 1 male and 4 female mice) in which we could evoke EPSCs from both pathways. Out of these neurons, activation of cNTS_cat_ afferents enhanced the amplitude of SpVc EPSCs in 6 (example in Fig 9A), reduced the amplitude in 1 and, no effect in 2. As a population, cNTS_cat_ stimulation increased the median EPSC_SpVc_ amplitude by 160 % (95% CIs: 100 to 200%), increasing from a baseline of 15 pA (95% CIs: 10 to 70 pA) to 22 pA (95% CIs: 10 to 120 pA, p = 0.016, Wilcoxon matched-pairs test, Figure 9B) following cNTS_cat_ stimulation.

In 6 neurons, we used paired pulse stimulation to examine whether cNTS_cat_ evoked changes in the amplitude EPSC_SpVc_ may arise from presynaptic mechanisms. Of these neurons, EPSC_SpVc_ was potentiated by cNTS_cat_ activation in 5 and depressed in 1. All 5 cells with potentiated EPSC_SpVc_ amplitudes also showed a decrease in PPR from a mean of 1.1 (95% CIs: 0.9 to 1.3) to 0.8 (95% CIs: 0.7 to 1.0), consistent with an increase in probability of release at these SpVc synapses. In the cell where cNTS_cat_ activation suppressed the amplitude of the evoked EPSC_SpVc_, the PPR increased from 0.8 to 0.9, consistent with a reduced probability of release. As a population, the mean PPR for all cells decreased from 1.0 (95% CIs: 0.8 to 1.3) to 0.8 (95% CIs: 0.7 to 1.0; paired t-test, p = 0.051, Fig. 9C). These data suggest that cNTS_cat_ afferents tend to increase synaptic activity in PB and potentiate release at SpVc synapses.

## Discussion

We show that noxious mechanical and thermal stimuli produce a sustained increase in norepinephrine (NE) in the parabrachial nucleus (PB), and that similar increases occur after direct stimulation of the cNTS, a primary source of NE input to PB via the A2 cell group. *In vitro*, photoactivation of catecholaminergic afferents from cNTS drives a prolonged increase in excitatory synaptic activity in parabrachial neurons. Activation of these afferents also potentiates synaptic transmission to parabrachial neurons from trigeminal sensory afferents. Together, these findings indicate that noradrenergic inputs from cNTS to PB may have long lasting faciliatory effects on nociception.

### Prolonged NE response to noxious stimuli

Brief noxious mechanical and thermal stimulation caused a sustained increase in NE levels in the parabrachial nucleus, measured with the fluorescent sensor NE2h. Contralaterally applied mechanical stimuli evoked larger transients than those applied ipsilaterally, while responses to ipsilateral and contralateral thermal stimuli were similar. Anatomically, projections from A2 primarily target the ipsilateral PB (Milner et al., 1986), consistent with the magnitude difference we observed following mechanical stimulation. The absence of a similar laterality effect with thermal stimulation may reflect a combination of experiment parameters (rapid mechanical vs gradual thermal stimuli) or may result from differences in how these modalities are detected and encoded. However, the similarity between contra- and ipsilateral evoked NE transients s suggests that the noradrenergic response reflects a general aversiveness of sensory inputs, rather than spatial or modality specific information. This functional observation is consistent with anatomical studies demonstrating bilateral projections from dorsal horn neurons to the NTS (Menetrey and Basbaum, 1987; Holt et al., 2019; Holt, 2022).

The prolonged signal recorded by the NE sensor is unlikely a reflection of the sensor kinetics. Medium and high affinity forms of this sensor (NE1m and NE1h) have off time constants shorter than 2 s (Feng et al., 2019), as does the second generation NE2h sensor used here (personal communication, Y. Li). This decay rate is substantially faster than our observed off-time constants of 44 s for thermal and 38 s for mechanical stimuli. Similarly prolonged NE responses to noxious or stressful stimuli have been measured using microdialysis. For example, noxious pin prick or cold stimuli produced prolonged elevations (up to 15 minutes) in NE levels the dorsal reticular nucleus of neuropathic rats (Martins et al., 2015) while intraplantar formalin injection produced a similarly prolonged elevation (∼15 minutes) of NE in the bed nucleus of the stria terminalis (BNST) (Deyama et al., 2008). Intermittent tails shocks cause prolonged elevations of NE in the prefrontal cortex and hippocampus of cold-stressed rats (∼ 30 to 60 minutes), (Nisenbaum et al., 1991; Gresch et al., 1994). Aversive stimuli also cause NE-dependent prolonged increases in firing rates of amygdala neurons (Giustino et al., 2020). Why these different stimulus modalities produce different transient profiles remains to be determined, however, these data indicate that comparatively brief noxious or stressful stimuli produce sustained NE signaling within the CNS, including in PB, as reported here.

### Source of NE inputs to parabrachial nucleus

Our focus on cNTS_cat_ neurons arises from a combination of anatomical, behavioral and electrophysiological evidence for their potential role in processing aversive stimuli. NE innervation of the parabrachial nucleus largely originates from the A2 cell group of the NTS, with a comparatively minor projection arising from the anatomically adjacent locus coeruleus (Milner et al., 1986). NTS neurons receive direct projections from medullary and dorsal horn neurons and solitary tract afferents, and this overlap has been postulated to aid integration of somatosensory and interoceptive stimuli (Menetrey and Basbaum, 1987; Appleyard et al., 2007; Holt et al., 2019; Holt, 2022). Projections from the cNTS, including A2 neurons, preferentially target the lateral regions of the parabrachial nucleus (Herbert et al., 1990), a hub for processing aversive stimuli (Palmiter, 2018; Uddin et al., 2018; Bowen et al., 2020; Raver et al., 2020). PB-projecting NTS neurons are activated by noxious peripheral stimuli (Figure 7) (Saito et al., 2017), and this activation is enhanced in a mouse model of trigeminal neuropathic pain (Okada et al., 2019).

A2 neurons respond to a wide range of stressors (Rinaman, 2011), including subcutaneous capsaicin or formalin, foot shocks, forced swim, and restraint stress (Pezzone et al., 1993; Palkovits et al., 1999; Rinaman, 2011). Following restraint stress, NE release from A2 afferents is potentiated in the bed nucleus of the stria terminalis (BNST) demonstrating that the output of these neurons is modified by aversive conditions (Schmidt et al., 2019). Our data suggests that A2 neurons also contribute to graded NE signaling in PB. Relatively brief stimulus trains in anesthetized mice evoked long lasting NE transients in PB that mirrored those evoked by noxious stimulation, and the magnitude of these transients was proportional to the stimulation frequency. We recognize that noxious and direct peripheral nerve stimulation also activates the locus coeruleus (Cedarbaum and Aghajanian, 1978; Guyenet and Byrum, 1985; Alba-Delgado et al., 2021) and projections from this cell group can regulate excitatory synapses in PB (Yang et al., 2021).

In addition to NE, the majority of A2 neurons co-express glutamate, as evidenced by mRNA expression for the vesicular glutamate transporter, VGLUT2, in 80% of A2 neurons (Stornetta et al., 2002) and by *in vitro* experiments where optical stimulation of A2 afferents evokes glutamatergic currents in parabrachial neurons (Roman et al., 2016). Although A2 afferent activation in this study appears to contain a similar glutamatergic component, additional experiments with specific antagonists are needed to determine the relative contributions of glutamatergic and adrenergic receptors to the PB neuron response profiles.

### NE increases excitability of parabrachial neurons

To facilitate NE release from cNTS_cat_ afferents, we used brief trains of stimuli known to drive NE release (Schmidt et al., 2019), and that resemble the firing profiles of A2 neurons *in vivo* (Moore and Guyenet, 1985). In neurons, in which the frequency of synaptic activity remained elevated for at least 20 s after the stimulus trai, this facilitation decreased with a time constant of 100 s (Fig 3E). This prolonged effect of cNTS_cat_ activation on synaptic activity *in vitro* is similar to the prolonged NE transients we observe in PB *in vivo* after noxious stimulation (Figs 1 & 2). Similar experiments targeting BNST demonstrate that cNTS_cat_ afferent activation can reliably produce prolonged NE transients *in vitro* (Schmidt et al., 2019). However, whether the facilitation of synaptic activity following cNTS_cat_ activation observed in our study result from a sustained NE transient remain to be determined.

Our data suggest that the prolonged effects of cNTS_cat_ stimulation on synaptic activity in PB primarily reflect presynaptic mechanisms. Paired pulse ratios (PPR) of optically evoked EPSCs from SpVc terminals were reduced by cNTS_cat_ afferent stimulation, a change thought to reflect an increase in the probability of transmitter release at presynaptic terminals (Thomson, 2000). In addition, the amplitude of the evoked SpVc afferent responses increased following cNTS_cat_ stimulation, despite a small reduction in membrane resistance. The shift in membrane resistance of PB neurons may reflect a postsynaptic effect of NE or may be a result of the increase in spontaneous synaptic activity produced by cNTS_cat_ stimulation.

## Conclusion

We show that activation of catecholaminergic afferents from cNTS increases excitatory network activity in PB and potentiates sensory inputs to this hub of nociception and aversion. As cNTS is a stress responsive hub for diverse interoceptive and exteroceptive inputs, this connection provides a pathway through which noxious events throughout the body may converge to increase the aversiveness of ongoing or new sources of pain. Understanding the mechanisms through which this occurs may help guide new therapies for treating chronic pain conditions.

## Acknowledgements

Research reported in this publication was supported by the University of Maryland Center to Advance Chronic Pain Research (CACPR) Collaborative Seed Grant Program and the National Institute of Neurological Disorders and Stroke of the National Institutes of Health grants R01NS099245 and R01NS069568. The content is solely the responsibility of the authors and does not necessarily represent the official views of the National Institutes of Health. The funding sources had no role in study design; the collection, analysis and interpretation of data; the writing of the report; or in the decision to submit the article for publication.

## Notes

### Competing Interest Statement

The authors have declared no competing interest.

### Summary of Updates

Fiber photometry data is now displayed as dF/F and data from individual mice are shown along with the group averages. Analysis of NE transient kinetics is updated to reflect differences in heat versus mechanical stimulation.

